# Rates of convergence in the two-island and isolation-with-migration models

**DOI:** 10.1101/2021.08.23.457399

**Authors:** Brandon Legried, Jonathan Terhorst

## Abstract

A number of powerful demographic inference methods have been developed in recent years, with the goal of fitting rich evolutionary models to genetic data obtained from many populations. In this paper we investigate the statistical performance of these methods in the specific case where there is continuous migration between populations. Compared with earlier work, migration significantly complicates the theoretical analysis and demands new techniques. We employ the theories of phase-type distributions and concentration of measure in order to study the two-island and isolation-with-migration models, resulting in both upper and lower bounds. For the upper bounds, we consider inferring rates of coalescent and migration on the basis of directly observing pairwise coalescent times, and, more realistically, when (conditionally) Poisson-distributed mutations dropped on latent trees are observed. We complement these upper bounds with information-theoretic lower bounds which establish a limit, in terms of sample size, below which inference is effectively impossible.

## 1 Introduction

Demographic inference–the estimation of past gene flow, migration, and size history events experienced by a population–is now a significant research area in evolutionary biology and mathematical genetics. Stimulated by an ongoing explosion in data availability, a series of increasingly sophisticated statistical methods has been developed to infer rich, highly parameterized demographic models using patterns of population genetic variation. These methods have seen significant uptake in biology, with the most successful (e.g., Gutenkunst et al., 2009; Li and Durbin, 2011) having been used in thousands of studies across a wide variety of species.

Formidable mathematical and computational hurdles must be overcome in order to estimate complex evolutionary models; often, even evaluating the likelihood function is nontrivial. As a result, research in this area has, to date, tended to focus on developing efficient inference methods. A much smaller number of authors have studied the question which interests us here: when is it theoretically (im)possible to estimate these models from data?

A starting point in the literature on the theoretical statistical aspects of demographic inference is Myers, Fefferman, and Patterson (2008), who proved the striking result that population size history is unidentifiable from the site frequency spectrum. That is, given an arbitrary size history function, there exists a smooth perturbation of it which produces exactly the same frequency spectrum in expectation. Subsequently, Bhaskar and Song (2014) showed that identifiability can be achieved by restricting the space of size history functions to be finite dimensional, for example piecewise constant or piecewise exponential. Terhorst and Song (2015) derived minimax lower bounds for demographic inference from the site frequency spectrum, and showed that there is a fundamental limit in our ability to infer size history for populations which have experienced a bottleneck. Baharian and Gravel (2018) showed that even in non-bottlenecked populations, there may be little to no statistical power to distinguish between different size history hypotheses on the basis of a finite amount of data. Working in a different setting, J. Kim et al. (2015) studied nonparametric estimation of the size history using samples of coalescent times from pairs of chromosomes, deriving both upper and lower bounds for hypothesis testing and estimation of the size history function. Johndrow and Palacios (2019) extended the analysis J. Kim et al. (2015) to coalescent trees on three samples, studied the benefit of incorporating ancient samples, and derived exact lower bounds on the Bayes error rate for distinguishing between population size histories.

All of the above papers consider the case of a panmictic population. Less attention still has been paid to inference in structured population models. Y. Kim et al. (2020) furthered the analysis of J. Kim et al. (2015) to the case where pairwise coalescent data is used to infer population structure, and showed in particular that the amount of data needed to accurately reconstruct the demography of a structured population may be exponential in the number of demes. Sousa, Grelaud, and Hey (2011) showed that the times of migration events in gene trees are not identifiable under a standard coalescent model.

A related thread concerns reconstructing the phylogeny or “species tree” of a set of populations tree under a structured coalescent model. Up to this point, coalescent-based approaches have mostly considered complications arising only from incomplete lineage sorting, which causes gene trees to have a different topology than the background species tree (Rannala and Yang, 2003; Allman, Degnan, and Rhodes, 2011; Mirarab et al., 2014). Although there has been some recent progress on phylogenetic inference with migration (Hey, Chung, et al., 2018; Flouri et al., 2019), the focus of this line of work, species tree estimation, is ultimately different from that of demographic inference, where we seek to infer distributional parameters over a collection of latent genealogies.

Despite these many interesting and useful contributions, it is fair to say that our ability to estimate complex demographic models has far outpaced our theoretical understanding of those estimators. As noted above, only a handful of theoretical studies consider inference in the presence of complex population structure. Nevertheless, such models are now routinely fit in practice, often using numerous populations and many different migration events (e.g., Gutenkunst et al., 2009; Jouganous et al., 2017; Kamm, Terhorst, and Song, 2017; Rodríguez et al., 2018; Kamm, Terhorst, Durbin, et al., 2020). Given that the theoretical results in the panmictic setting have so far been mainly negative, it seems important to extend the analysis to other types of structured population models that are becoming prevalent.

In this paper, we address this gap by theoretically analyzing some inference problems that arise in a structured coalescent model with continuous migration. Although some of our proof techniques are based on these earlier works (in particular, that of J. Kim et al., 2015), as we will see, migration introduces significant challenges into the analysis, requiring different approaches than have been used previously. Consequently, we restrict our focus to the simplest non-trivial structured population model of two islands with continuous migration between them, and a variant of it known as the isolation-with-migration model. In Section 2, we lay out our notation. Section 3 formalizes the model and introduces key definitions. Section 4 derives some eigenvalue and tail bounds for the two-island model, some of which may be useful more generally. Section 5 studies moment-based estimation of the key parameters in the two-island model. Section 6 derives information-theoretic lower bounds on the ability to distinguish different island models from data. Section 7 concludes with some discussion.

## 2 Notation

Throughout the paper, *n* is used to denote sample size, and we suppress explicit dependence on it when there is no possibility of confusion. For any *r* > 0 and x ∈ ℝ^*d*^, let *B*_*r*_(x) be the ball of radius *r* around x, i.e.

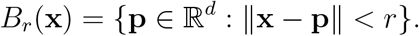

The constant

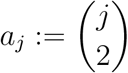

appears throughout the paper.

Matrices and vectors are denoted in boldface. The *L*^*p*^ norm of x is denoted

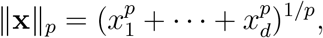

and the *L*^∞^ norm is denoted ‖x‖_∞_ = max_*i*_ |*x*_*i*_|. If *p* is not indicated, then 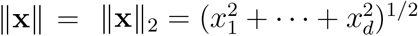 is taken to be the *L*^2^ (Euclidean) norm. If **A** ∈ ℝ^*m*×*n*^ is a matrix, then

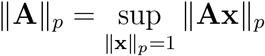

denotes the induced *p*-norm, with ‖**A**‖ denoting the operator norm. The Frobenius norm is given by

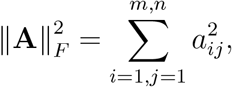

where *a*_*ij*_ is the *ij*th entry of the matrix **A**. If *m* = *n* such that **A** is square, then the trace and determinant of **A** are denoted tr **A** and det **A**, respectively. The identity matrix is denoted **I**. The standard basis vectors are denoted

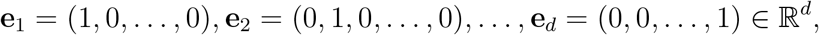

and the vector of all ones is denoted **e** = (1, 1, …, 1)^**T**^. The dimensionality of **I, e**_*i*_, and **e** may vary from usage to usage, but will be obvious from context.

In this paper, the dimension of all vector spaces is either 3 or 4, independent of any other problem-specific quantities. Hence, by equivalence of matrix norms, there exist universal constants *C*_1_, *C*_2_ > 0 such that

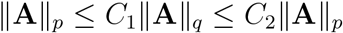

for all *p, q* > 0, including *p* = ∞ or *p* = *F* (Frobenius norm). In particular, for **A** ∈ ℝ^4^ we have

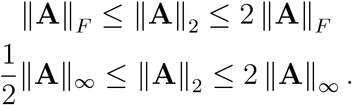

## 3 The model

We consider a structured coalescent model with two demes and continuous migration between them, sometimes referred to as the “two-island” model (Takahata, 1988; Notohara, 1990). The model considers the probability distribution of a genealogy formed by sampling a pair of chromosomes. The time *t* = 0 corresponds to the present while positive *t* corresponds to *t* generations in the past. For any *t* ≥ 0, the vector

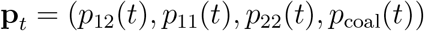

gives a probability distribution on the finite sample space Ω = {12, 11, 22, coal}. Here, *p*_12_(*t*) is the probability that a pair of individuals sampled in the present descend from a pair of individuals separated into the islands 1 and 2 at time *t*. Similarly, *p*_11_(*t*) is the probability that they descend from a pair of individuals both on island 1 at time *t*, with an analogous definition for *p*_22_(*t*). Lastly, *p*_coal_(*t*) is the probability that they descend from a common ancestor at time *t*.

Let *m*_1_ be the rate at which an individual migrates from island 1 to island 2, similarly for *m*_2_. For any pair of individuals in the present, the time to their most recent common ancestor is called the coalescent time. Let *c*_1_ be the corresponding rate of coalescence for the two individuals if they both live on island 1 and *c*_2_ be the respective rate for island 2. The movement of a pair of individuals between these four states is modeled by a continuous-time Markov chain (CTMC) with state probabilities

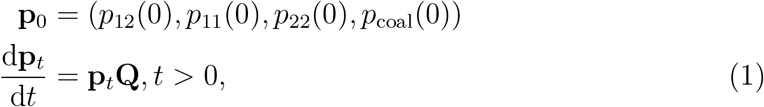

where **Q** is the transition rate matrix

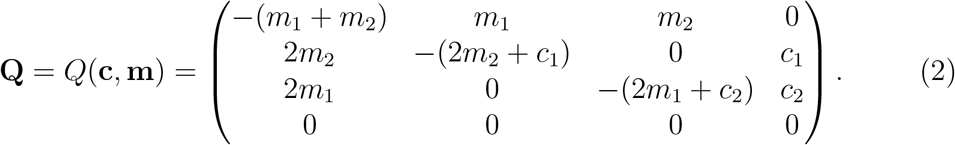

To avoid degeneracies, we assume henceforth that both of the coalescent rates *c*_*i*_ and at least one of the migration rates *m*_*i*_ are strictly positive. At the present *t* = 0, we sample two individuals assuming they are not coalesced, so *p*_coal_(0) = 0. The solution to (1) is **p**_*t*_ = **p**_0_*e*^**Q***t*^.

For *x* ≥ 0 and *t* > *x*, the hazard rate of coalescence *h*(*t* | *x*) = *h*^**c**,**m**^(*t* | *x*) is the rate of entry into the “coal” state, given that the process has not already done so up to time *x*. Viewing *t* = *x* as the present time, conditioning on noncoalescence implies *p*_coal_(*x*|*x*) = 0. If we define **p**_*t*|*x*_ to be the distribution of the process at time *t* ≥ *x* conditioned on noncoalescence up to time *x*, then

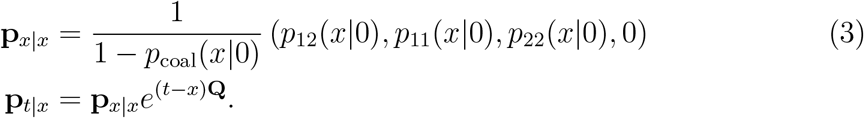

The (conditional) hazard rate of coalescence at time *t* is given by multiplying **p**_*t*|*x*_ with the fourth column of **Q**:

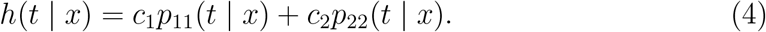

Suppose *n* ≥ 2 individuals are sampled at time *t* = 0 and consider the sequence of coalescent times 0 = *x*_*n*+1_ < *x*_*n*_ < … < *x*_2_ in the genealogy of the sample. Any pair of non-coalesced lineages is as likely to coalesce as any other, so the conditional hazard rate for coalescent time *x* is then given by

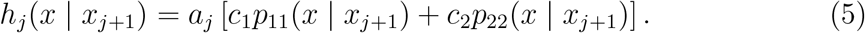

The conditional density of the *j*-th coalescent time given the (*j* + 1)-th coalescent time is

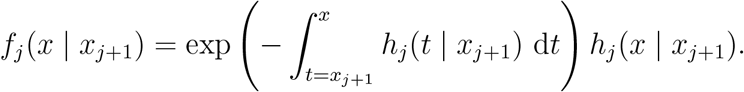

From the Markov property, the joint density of the coalescent times *x*_*n*_ < … *< x*_2_ is then

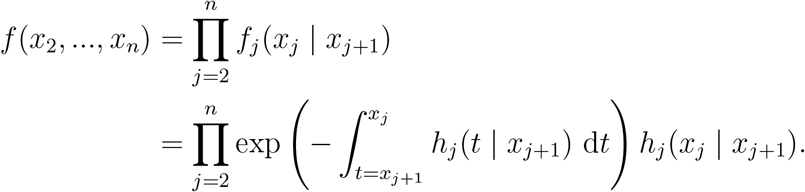

## 4 Tail bounds

We now collect some results about the two-island model with migration which will be used below. These results exploit the fact that coalescent times in this model follow a so-called phase-type distribution. A useful reference on these distributions is Asmussen and Albrecher (2010). Hobolth, Siri-Jegousse, and Bladt (2019) have also recently studied phase-type distributions in a related setting.

### Definition 1

(Phase-type distribution). Let {*X*_*t*_} _*t*≥0_ be a homogeneous, continuous time Markov chain on a finite state space 𝒮 with single absorbing state Δ ∈ 𝒮. The rate matrix of *X*_*t*_ may be written in block form as

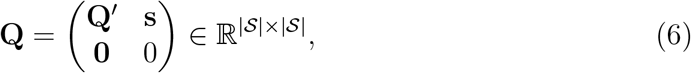

where **Q**′ ∈ ℝ^(|𝒮|−1)×(|𝒮|−1)^ is the sub-intensity matrix giving the transition rates between the transient states in 𝒮, and **s** ∈ ℝ^|𝒮|−1^ is a column vector giving the transition rates from each transient state to Δ. Also, let ***α*** ∈ ℝ^|𝒮|^ be the distribution of *X*_0_. The first hitting time *ζ* = inf {*t* > 0 : *X*_*t*_ = Δ} is said to be of *phase-type with representation* (**Q, *α***). We denote this as

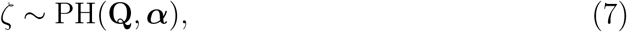

and denote the distribution function of *ζ* by ℙ_***α***_(*ζ < t*).

Below we will need to simultaneously consider phase-type distributions with multiple initial distributions ***α***_*ij*_, *i, j* ∈ {1, 2*}*. We slightly abuse notation and denote these by 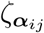.

An important quantity in the theory of phase-type distributions is the spectrum of **Q**. In particular, the long-term behavior of the process is controlled by the size of the gap between the largest and second-largest eigenvalues. Our first result quantifies this gap for the chain defined in (2).

### Proposition 2.

*Let* **Q** = *Q*(**c, m**). *Then the eigenvalues λ*_0_, …, *λ*_3_ *of* **Q** *are non-positive and real: λ*_3_ ≤ *λ*_2_ ≤ *λ*_1_ *< λ*_0_ = 0, *with*

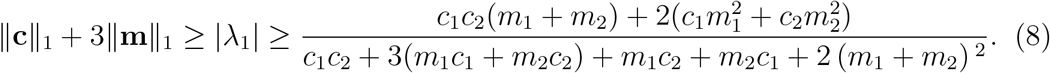

*Proof*. By the Gershgorin circle theorem, the eigenvalues of **Q** have nonpositive real part, and *λ*_0_ = 0 since **Q** is a rate matrix. Let **Q**′ be the 3 × 3 leading principal minor of **Q**. Then *v* = (*v*_1_, *v*_2_, *v*_3_, *v*_4_)^**T**^ is an eigenvector of **Q** with nonzero eigenvalue if and only if *v*_4_ = 0 and (*v*_1_, *v*_2_, *v*_3_)^**T**^ is an eigenvector of **Q**′ with the same eigenvalue.

Defining 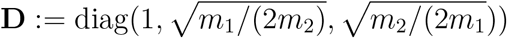, we have

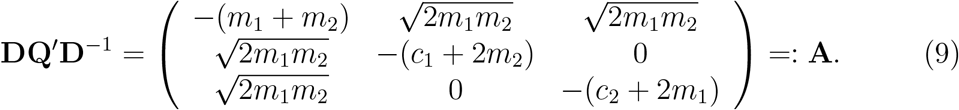

Thus, **Q**′ is similar to the Hermitian matrix **A**, so *{λ*_1_, *λ*_2_, *λ*_3_*} ⊂* ℝ_≤0_. In fact, since

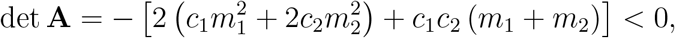

**A** has no zero eigenvalues: *λ*_3_ ≤ *λ*_2_ ≤ *λ*_1_ < 0. Finally,

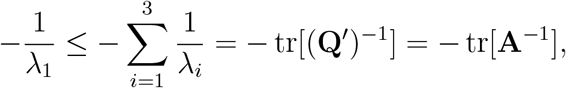

which yields the lower bound in (8) by direct computation of **A**^−1^. Finally, for the upper bound we have

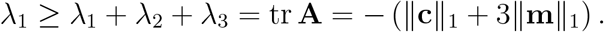

□

Since only the leading eigenvalue plays a role in the sequel, we define

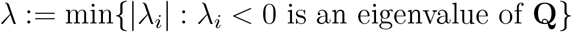

for the rest of the paper.

The following quantity appears repeatedly in the results to come.

### Definition 3.

Given a rate matrix **Q** with leading eigenvalue *λ*, the *condition number* of **Q** is *κ* := ‖**Q**‖*/λ*.

*Remark. κ* differs slightly from the usual definition: it is the ratio of the largest singular value to smallest (absolute) eigenvalue of **Q**.

### Lemma 4.

*Let* 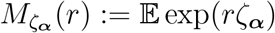 *be the moment generating function of ζ in* (7). *Then* 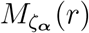 *is defined for all r < λ. Furthermore, for any such r*,

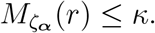

*Proof*. Let *λ*_3_ ≤ *λ*_2_ ≤ *λ*_1_ < 0 be defined as in Proposition 2. By Proposition IX.1.8 of Asmussen and Albrecher (2010), 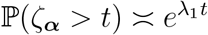 as *t →* ∞, which implies that

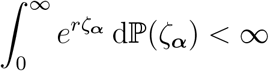

for *r <* |*λ*_1_|. Next, by Proposition IX.1.7 of Asmussen and Albrecher,

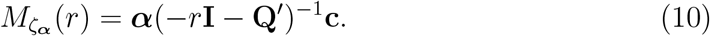

With **Λ** = diag(*λ*_1_, *λ*_2_, *λ*_3_) and 0 < *r* < |*λ*_1_|, we get

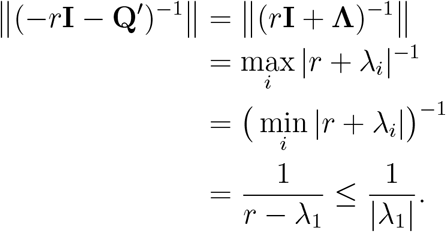

The result follows from (10) and the facts that

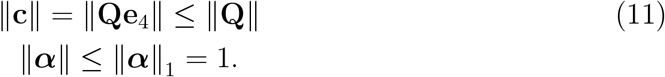

□

### Corollary 5.

*For any t* > 0,

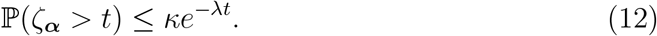

*Proof*. Apply the previous lemma to the Chernoff-type bound

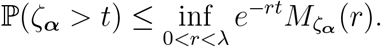

□

The final result of this section establishes that the two-island model forgets its starting state exponentially quickly.

### Proposition 6.

*For any two initial distributions* ***α***_1_, ***α***_2_, *we have*

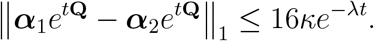

*Proof*. Let ℙ_***α***_(*X*(*t*) = *s*) be the *s*th component of ***α****e*^*t***Q**^. For *i* = 1, 2 define

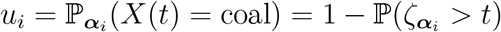

for 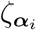 defined by (7). Then for *s* ≠ coal,

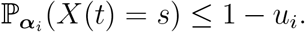

By repeated applications of the triangle inequality,

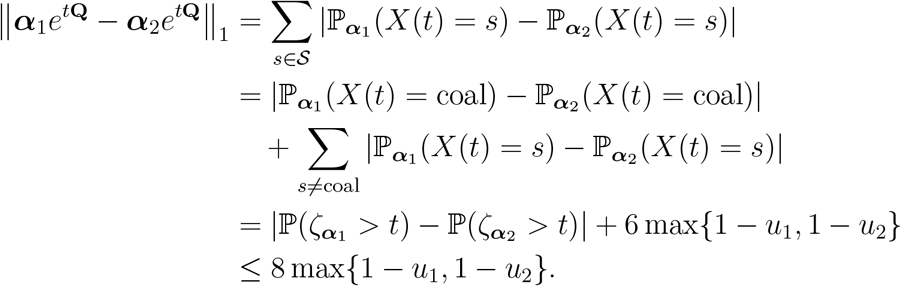

Finally, by Corollary 5,

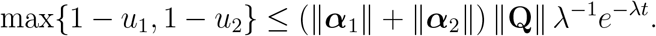

□

The next results pertain to two possibly different island models specified by rate matrices **Q**^(*i*)^ = *Q*(**c**^(*i*)^, **m**^(*i*)^) (cf. equation 2) for *i* = 1, 2, with leading eigenvalues *λ*^(*i*)^ and condition numbers *κ*^(*i*)^. We let 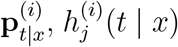, and *f* ^(*i*)^ refer to the transition probability, hazard rate, and joint density for the corresponding model.

### Definition 7.

Given a pair of two-island models **Q**^(1)^ = *Q*(**c**^(1)^, **m**^(1)^) and **Q**^(2)^ = *Q*(**c**^(2)^, **m**^(2)^), we say *model 2 is δ-close to model 1* if there exist diagonal matrices **D**_*c*_, **D**_*m*_ with max{‖**D**_*c*_‖, ‖**D**_*m*_‖} < *δ* such that

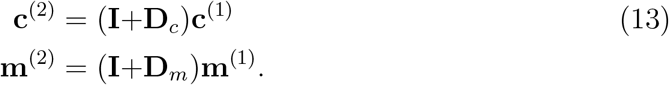

By abuse of notation, we also refer to **Q**^(2)^ being *δ*-close to **Q**^(1)^. It follows from the definition that if **Q**^(2)^ is *δ*-close to **Q**^(1)^ then

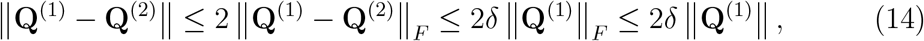

whence

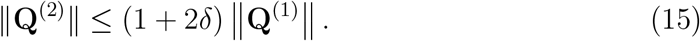

### Lemma 8.

*If* **Q**^(2)^ *is δ-close to* **Q**^(1)^ *then*

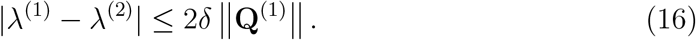

*Proof*. As in the proof of Proposition 2, let **A**^(1)^ and **A**^(2)^ be the Hermitian matrices to which (the upper-left submatrices of) **Q**^(1)^ and **Q**^(2)^ are similar, cf. equation (9). By Weyl’s eigenvalue perturbation theorem (e.g., Horn and Johnson, 2012, Theorem 4.3.1)

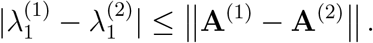

By the stated assumptions, we have for the entries of **A**^(1)^ and **A**^(2)^,

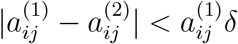

which implies that

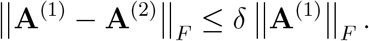

Hence,

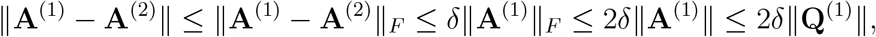

where the final inequality is because

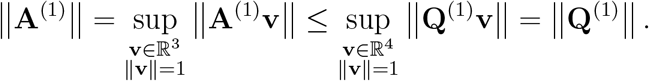

□

### Proposition 9.

*Suppose* **Q**^(1)^ *and* **Q**^(2)^ *are δ-close for some δ <* 1/(4*κ*^(1)^). *Then for all t* ≥ 0,

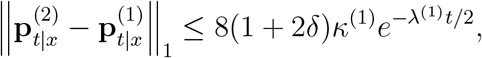

*Proof*. By Lemma 8 and the assumptions,

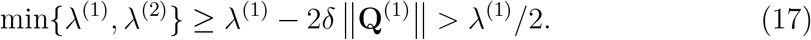

Observing that **e**_4_ = (0, 0, 0, 1)^**T**^ is a left eigenvector of *e*^*t***Q**^ for any *t* and **Q** defined by (2), we have

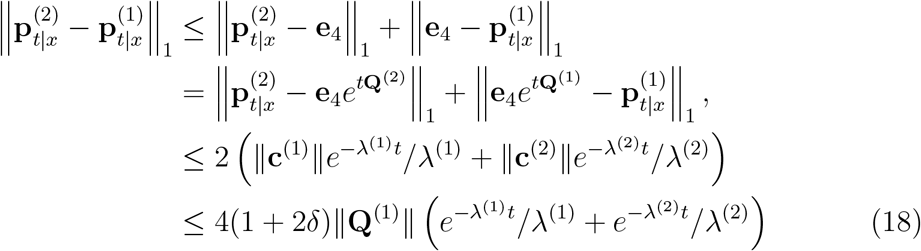

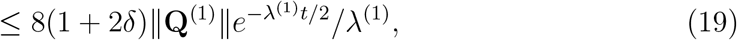

where inequality (18) follows from equations (11) and (15), and inequality (19) follows from (17) and the fact that *x* ↦ *e*^−*xt*^/*x* is decreasing.

□

The second bound establishes convergence to zero linearly in *δ*. This result is due to Mitrophanov (2003, Corollary 2.1); we state it in an adapted form below.

### Theorem 10

(Mitrophanov 2003). *Let* **Q**^(1)^ *and* **Q**^(2)^ *be as in Proposition 9, and further suppose that*

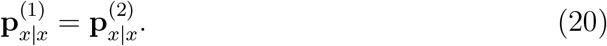

*Then*

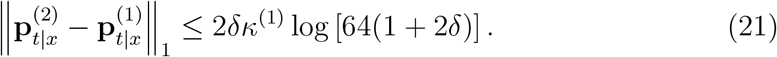

*Proof*. In the notation of Mitrophanov (2003), Proposition 6 implies *b* = *λ*^(1)^/2, *c* = 64(1 + 2*δ*) > 2 in his equation (2.1), and (20) implies **z**(*t*) = 0. Plugging these constants into Mitrophanov’s equation (2.9) and using (14), we obtain

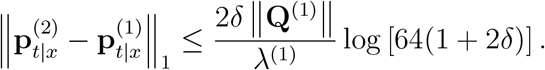

□

## 5 Upper bounds

In this section, we derive an estimator for the parameters in the symmetric two-island model, and prove some results about its accuracy on finite samples. Before presenting our results, we first outline some of the challenges of inference in structured population models.

In the panmictic setting, J. Kim et al. (2015) derive a nonparametric estimator of the effective population size function *N*(*t*) based on some ideas from survival analysis. They rely on an explicit expression for the hazard rate function ℙ(*T*∈ [*x, x* + *dx*) |*T* > *x, N* (*t*)) of the coalescent time *T* (see Remark 2.1 in their paper). This function is then inverted, yielding a histogram-type estimator for 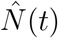. The simple form of the estimator makes it possible to precisely analyze its performance on finite samples.

Unfortunately, it does not seem possible to extend their approach to the case of (even relatively simple) structured population models. The hazard rate function (4) depends on **m** and **c** in a complicated way, and cannot be analytically inverted. Thus, although classical (asymptotic) guarantees are obtainable, it is difficult to study the finite-sample behavior of likelihood-based estimators in structured population models.

To make progress, we turn to a moment-based estimator instead. Let *E*_12_ be the expected time to coalescence for a pair of individuals that live on different islands and *E*_11_, *E*_22_ be the analogous expectations for pairs living on the same islands. With **Q**′ defined as in Section 3, we have

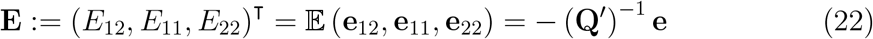

where the initial distribution **e**_*ij*_ places all mass on state *ij*, and the final equality is a property of phase-type distributions (Asmussen and Albrecher, 2010, Theorem IX.1.5).

Three parameters may be estimated from the first moments. We thus restrict our attention to the symmetric two-island model where *m*_1_ = *m*_2_ = *m*, wherefore

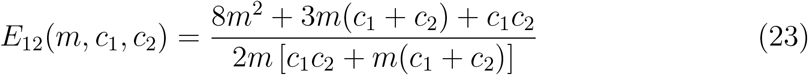

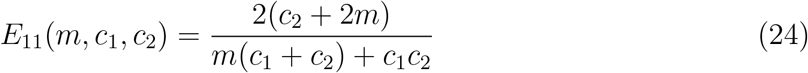

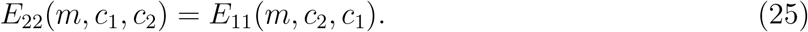

*Remark*. Note that the expected time to coalescence for two lineages sampled from the same deme is invariant to *m* when *c*_1_ = *c*_2_, a special case of Strobeck’s theorem (Strobeck, 1987; Durrett, 2008).

For each *ij* ∈ {12, 11, 22*}*, suppose we sample *L* i.i.d. coalescent times 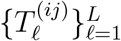 and form the sample version **Ê** of **E** by averaging:

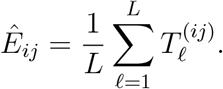

By the law of large numbers, **Ê** is a consistent estimator of **E**. Given **Ê**, we solve (23)–(25) for *m, c*_1_, *c*_2_ to obtain the following consistent estimators of the model parameters:

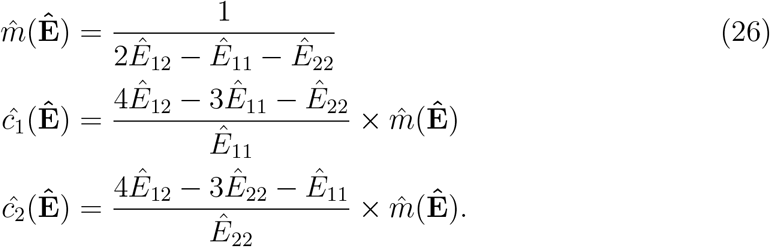

### 5.1 Error analysis

Now we derive bounds on the estimation error as a function of the number of samples *L*. Noting that the numerator and denominator of 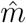 and *ĉ*_1_ (we omit discussion of *ĉ*_2_ since it is symmetric to *ĉ*_1_) are both homogeneous polynomials in **Ê**, this is most easily accomplished by considering the relative error.

#### Proposition 11.

*Suppose that* |*E*_*ij*_ *− Ê*_*ij*_ | ≤ *δE*_*ij*_ *for i, j* ∈ {1, 2*}. Then*

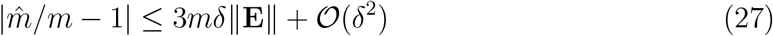

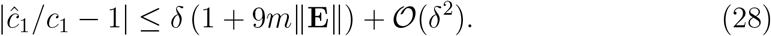

*Proof*. The supposition is equivalent to **Ê** =(**I** + **D**)**E**, where **D** is a diagonal matrix with ‖**D**‖ ≤ *δ*. Thus, with **a**_*m*_ = (2, −1, −1)^**T**^, we get

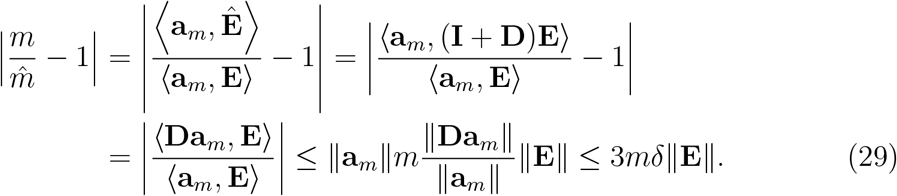

Equation (27) follows since 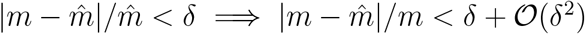.

Similarly, for *ĉ*_1_ and **a**_*c*_ = (4, −3, −1)^**T**^,

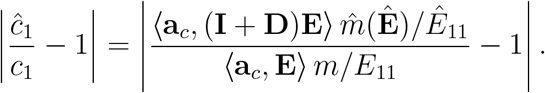

By our assumptions and the relative error bound (29), there exist *ϵ*_1_, *ϵ*_2_ such that

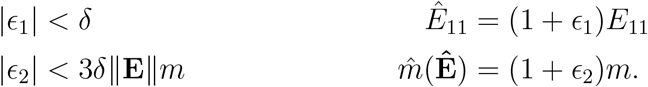

Then

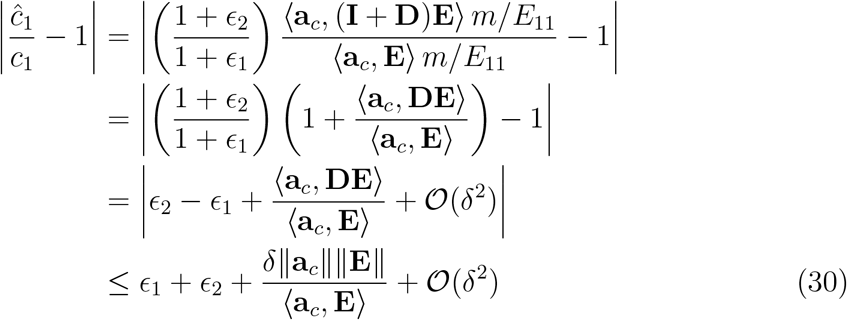

Now since ‖**a**_*c*_‖ < 6 and

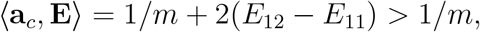

we have

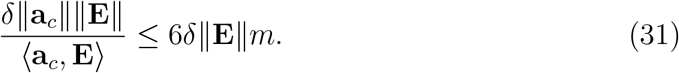

Inserting (31) into (30) and simplifying yields (28).

□

Next, we show that **Ê** is concentrated around its expectation **E**. This essentially follows from the fact that **Ê** is the sample average of phase-type distributions (see equation 22), and the tail bounds we derived in Section 3.

#### Proposition 12.

*Let* 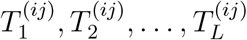 *be i*.*i*.*d. with distribution* PH(**Q, *α***_*ij*_). *Then*

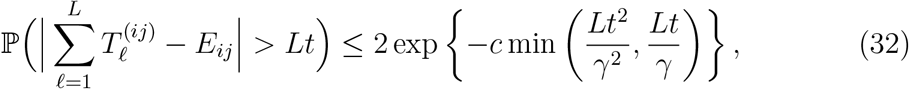

*where*

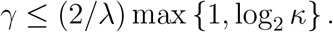

*and c* > 0 *is a universal constant*.

*Proof*. By Jensen’s inequality and Lemma 4, for sufficiently small *r* and any *K* ≥ 1,

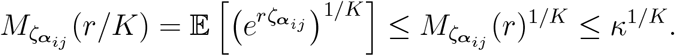

Let

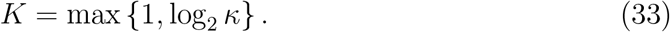

Then 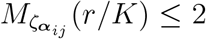. This implies that 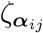 has a *sub-exponential* distribution, in the sense of Vershynin (2018, Proposition 2.7.1), with (see Vershynin, 2018, Definition 2.7.5)

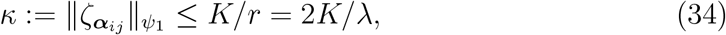

where 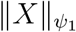 denotes the Orlicz 1-norm of the random variable *X*, and we chose *r* = *λ/*2 (say). The bound (32) then follows from Bernstein’s inequality (Vershynin, 2018, Corollary 2.8.3).

□

As we have seen, the leading eigenvalue *λ* factors integrally into our convergence rates. To gain intuition, consider the completely symmetric case where *m*_1_ = *m*_2_ = *m* and *c*_1_ = *c*_2_ = *c*. Then by (8),

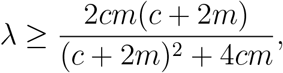

and we can distinguish a few cases:

- If *m* ≪ *c* then *λ* is roughly lower-bounded by 2*m*. This occurs when there is a high rate of migration between two islands with small effective population sizes. We then have log (‖**c**‖/*λ*) ≈ log [‖**c**‖/(2*m*)] ≫1. The bound (34) degenerates, such that we cannot rule out *κ* ≫ 1. In turn, the concentration inequality (32) degrades and we longer have good control on ‖**E** − **Ê**‖. This result quantifies the intuitive statement that inference (in particular, estimation of *m*) is difficult when the rate of migration is small.
- If *m* ≫ *c*, then *λ* ≈ *c*/2. This occurs when there is migration between two islands with large effective population sizes. Then 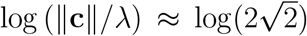, so *κ* ∈ 𝒪 (*c*) in equation (34). The right-hand side of (32) is essentially exp(*−Lt/c*), and the rate of convergence is dominated by the overall rate of coalescence.

### 5.2 Poisson hierarchical model

The previous section derives error bounds under the assumption that we could directly sample pairwise coalescent times within and between demes. In this section, we relax this unrealistically favorable assumption, and instead consider a model where the data consist of counts of the number of pairwise mismatches between randomly sampled genes. Specifically, we suppose

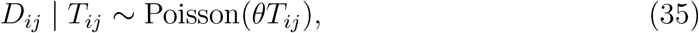

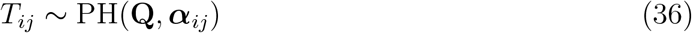

where the initial distribution ***α***_*ij*_ places all mass on deme *ij* ∈ {11, 22, 12}, so that *T*_*ij*_ are (i.i.d.) pairwise coalescent times between two genes sampled from demes *i* and *j*, which may be equal. Here *θ/*2 is the rate of mutation per unit of coalescent time, assumed known. This is a realistic data generating model if there is no recombination within genes; free recombination between genes; and *θ* is low.

For each *ij* ∈ {11, 22, 12*}*, we sample *L* i.i.d. mutation counts 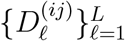 and estimate the *E*_*k*_ using sample averages:

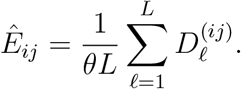

Then we use the estimators developed in the previous section.

To get a rate of convergence, we need to extend Proposition 12 to the marginal distribution of *D*_*ij*_ in (35).

#### Lemma 13.

*Let D*_*ij*_ *be distributed according to* (35)*–*(36), *and let* 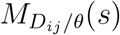 *denote the moment generating function of D*_*ij*_ */θ. Then for all s* ≤ *θ* log(1 + *λ/θ*),

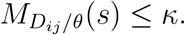

*Proof*. We have

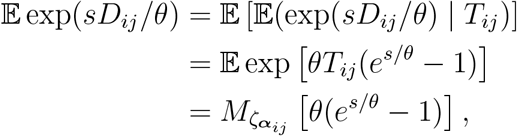

so the claim follows from Lemma 4.

□

*Remark*. Lemma 13 also follows from general results on subexponential mixtures of Poisson random variables (Schmidli, 1999), but our earlier results enable a direct and more quantitative proof.

#### Proposition 14.

*Let* 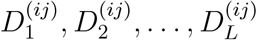 *be i*.*i*.*d. with distribution D*_*ij*_ *in* (35). *Then*

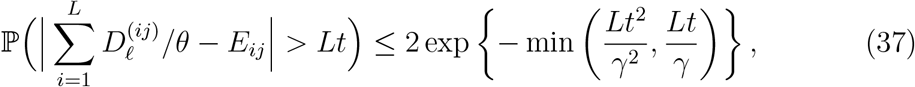

*where*

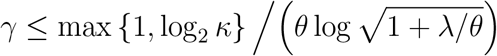

*and c* > 0 *is a universal constant*.

*Proof*. As in the proof of Proposition 12, we find that 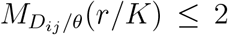 for *r < θ* log(1 + *λ/θ*) and the same constant *K*. Taking 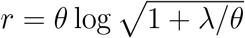, weget

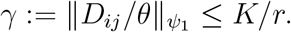

□

### 5.3 Simulations

Using Propositions 11–13, we can bound the accuracy of migration and coalescent rate estimates in the two-island model from finite amounts of data. For example, setting *t* = *δE*_*ij*_ in (37) and finding *L* such that the right-hand s ide is less than or equal to *ϵ*, we get a bound on the relative error |*Ê*_*ij*_ *− E*_*ij*_ |*/E*_*ij*_ *< δ* that holds with probability at least 1 *− ϵ*.

In Figure 1 we consider using sample averages of *T*_*ij*_ and *D*_*ij*_ to estimate *Ê*_12_, the average coalescent time for lineages originating in different demes. We set *δ* = *ϵ* = 0.1, i.e. < 10% relative error with > 90% probability, and for simplicity we took *c*_1_ = *c*_2_ = 1. In the simulations of *D*_*ij*_, the mutation rate was set to *θ* = 0.1. The area between the shaded blue region contains the .05–.95 quantiles of the sampling distribution of |*Ê*_12_ *− E*_12_| */E*_12_, obtained over 1,000 independent trials. The red lines are (1 ±0.1)*E*_12_. Based on our theoretical calculations, we found the value of *L*\* needed to ensure that the blue region was contained between the red lines. Thus, the sharpness of our bounds is reflected in the gap between the red lines (bounds) and blue region, with a larger gap indicating that our bounds predicted more samples were required than were actually necessary.

**Figure 1:**
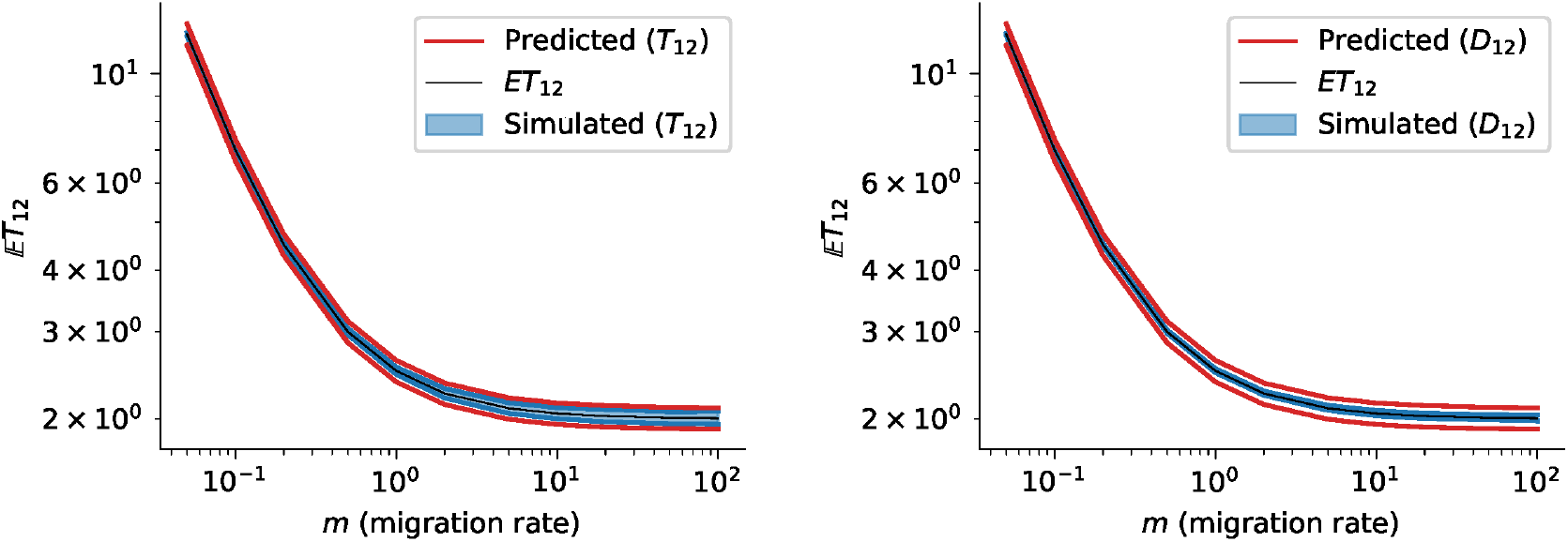
Observed and predicted 90% confidence intervals for the 5^th^ and 95^th^ percentiles of the sample mean. Left panel: directly sampling (*T*_*ij*_). Right panel: Poisson noise (*D*_*ij*_).

We see that the bounds are fairly accurate, particularly for using *T*_12_ (direct sampling of coalescent times) in order to estimate the population means. The actual number of samples *L*^*∗*^ is plotted in Figure 2 (left panel). As expected, estimating *E*_12_ with Poisson noise is more difficult, requiring 1–2 order of magnitude more data to obtain the same level of accuracy, and estimation requires more data when the migration rate is low.

**Figure 2:**
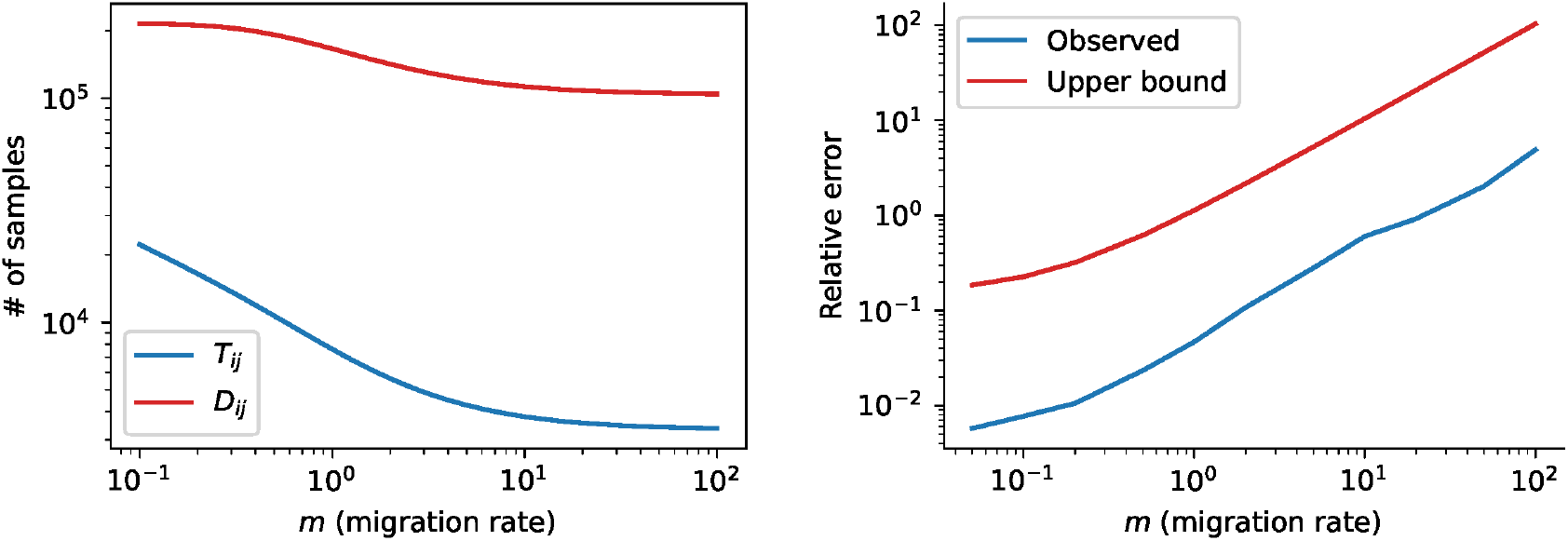
(Left panel) Number of samples calculated to obtain concentration bounds in Figure 1. (Right panel) Relative error in estimating 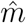 from Poisson-distributed mutation data.

Next, we used simulations of *D*_*ij*_ to estimate the migration rate *m* using (26). We studied the relative error 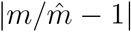 and compared it to the bound (27), where, as noted above, *δ* = 0.1 and *m* varied across a range of values. (Of course, (29) depends on the true parameters *m* and ‖**E**‖, so the bound may have limited practical use, but we can use it to get intuition for how the methods perform on real data.) The results are shown in the right panel of Figure 2, where we plugged *δ* = 0.1 and relevant values *c*_1_, *c*_2_, *m* and **E** into (29) to obtain the upper bound. We can see that the bound is loose by a (large) constant, but has the correct functional dependence in *m*. Since Figure 1 showed that the concentration bounds on *D*_*ij*_ are accurate, this imprecision is probably due to the fairly rudimentary bounds employed in the proof of Proposition 11.

## 6 Lower bounds

In this section, we prove several lower bounds on parameter estimation in the two-island migration models. The starting point of our work is the following result of J. Kim et al. (2015) on distinguishing between different single-population coalescent models.

**Theorem** (J. Kim et al. 2015, Theorem 3.2). Consider the following hypothesis testing problem: *H*_1_ states that the effective population size during the interval [0, ∞) is constant *N*, while *H*_2_ states that the population size during the same interval is the constant (1 + *η*)*N* for a fixed *η* > 0. If *L* i.i.d. coalescent trees on *n* individuals are observed from either *H*_1_ or *H*_2_, each with prior probability 1/2, then the Bayes error rate for any classifier is at least (1 − ϒ)/2, where

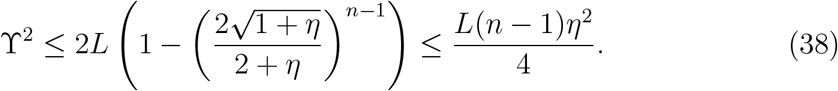

In this section, we prove analogous results for population sizes and migration parameters in the two-island model and the isolation-with-migration model. There is a limitation in distinguishing between two hypotheses on population history for an arbitrary period of time with any estimation method, even though simple moment-based estimators converge quickly to their respective model parameters.

### 6.1 Probability metrics and Bayes error rate

The section expands on the discussion in Section 3 of J. Kim et al. (2015). Let (Ω, ℱ, *P*) and (Ω, ℱ, *Q*) be two measures defined on a common probability space, with corresponding probability density functions *f*_*P*_ and *f*_*Q*_. The *total variation distance* between *P* and *Q* is defined to be

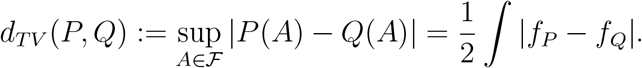

By abuse of notation, we may sometimes write *d*_*TV*_ (*f*_*P*_, *f*_*Q*_) to mean the same thing.

Suppose we are given a datum *D* that has been generated under either *P* or *Q*, and are asked to decide which measure was used assuming both choices are equally likely. The total variation distance between *P* and *Q* bounds the ability of any classifier to do so. Indeed, let *χ* ∈ *{P, Q}* denote the true data generating distribution, and 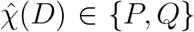 be a classifier. The probability that 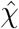 correctly classifies *D* can be written

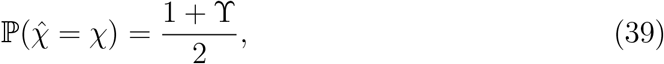

where ϒ > 0 since the error of any binary classifier can be made less than 1 /2. Note that (39) rearranges to

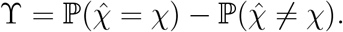

It can be shown (e.g., Devroye, Györfi, a nd Lugosi, 2 013) that the best possible classification rule is the likelihood ratio: is 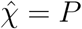 iff *P* (*D*) > *Q*(*D*), in which case

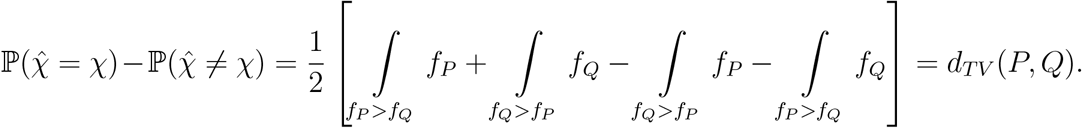

This classification rule is said to achieve the minimal or “Bayes” error rate. If multiple samples are given, say *D*_1_, …, *D*_*L*_, then ϒ = *d*_*T V*_ (*P*^⊗*L*^, *Q*^⊗*L*^), where *P*^⊗*L*^ denotes product measure.

In our setting, it is easier to work with a related quantity know n as the Hellinger distance:

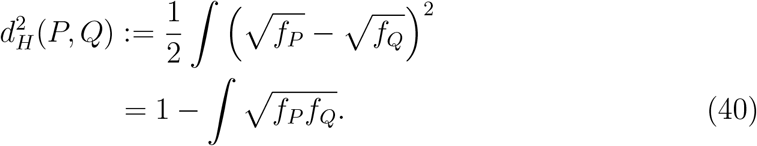

It is easily shown that 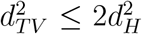. Furthermore, the Hellinger distance distributes over product measures: if *P* = *P*_1_ ×*P*_2_ and *Q* = *Q*_1_ ×*Q*_2_ represent product measures, then

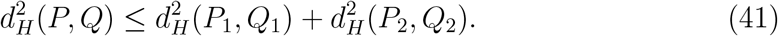

Hence, given *L* i.i.d. samples hypothesized to have been generated under either *P* or *Q*, it follows that

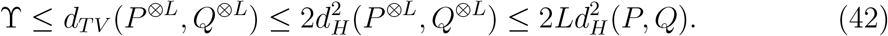

Going forward, we may abuse notation by identifying ℚ_*i*_ with its corresponding probability density function *f* ^(*i*)^ and compute 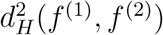 and *d*_*TV*_ (*f* ^(1)^, *f* ^(2)^).

### 6.2 Two-island models

In this subsection, we study the ability to statistically distinguish between different two-island models as a function of how close they are to each other. For *i* = 1, 2, we suppose that under *H*_*i*_ the coalescent times are generated under a two-island model with rate matrix **Q**^(*i*)^ = *Q*(**c**^(*i*)^, **m**^(*i*)^), and that **Q**^(2)^ is close to **Q**^(1)^ in a sense that is made precise below.

To improve readability, for the remainder of the section we suppress dependence of the density and hazard functions on *x* and *y* when there is no risk of confusion.

Let 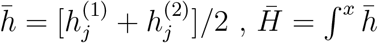, and 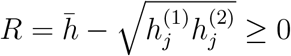. Then

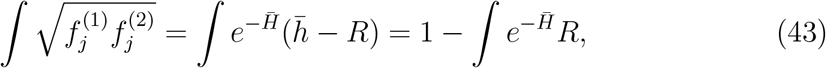

all integrals being over the positive reals. So by (40),

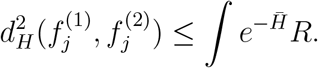

If 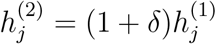 then 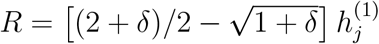, whence

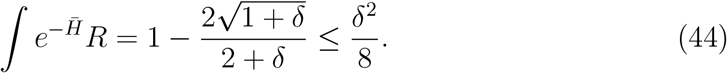

This is essentially the bound obtained by J. Kim et al. (2015, Theorem 3.2).

Below we extend this result to the two-island model. Theorem 15 covers the case when the two model hypothesis are *δ*-close in the sense of Definition 7, without placing any additional assumptions on the relationship between the hypotheses. This result is general, but as can be seen from equation (45), the bound is on the order 𝒪(*δ*), so it is asymptotically looser than the 𝒪(*δ*^2^) bound indicated by (44). Getting the 𝒪(*δ*^2^)rate turns out to depend rather delicately on a cancellation of the first-order coefficients in the Taylor expansion of 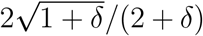. This, in turn, only seems to happen if the hazard rate function 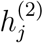 is an exact scalar multiple of 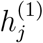. When there is more than population, such an equality no longer holds *even when* the parameters of the underlying model differ only by a multiplicative factor. Currently, we do not know if the difference in rates is an artifact of our proof technique, or if having data from multiple populations in fact renders the inference problem quantitatively easier.

#### Theorem 15.

*Let H*_1_ *and H*_2_ *be hypotheses with corresponding rate matrices* **Q**^(1)^ *and* **Q**^(2)^ *such that H*_2_ *is δ-close to H*_1_. *If L i*.*i*.*d. pairwise coalescent times are sampled under H*_1_ *or H*_2_, *each with probability* 1/2, *then for sufficiently small δ* > 0, *the Bayes error rate for any classifier is at least* (1 − ϒ)/2, *where*

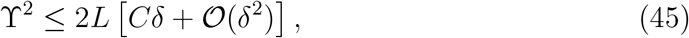

*where*

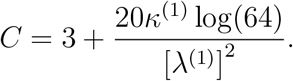

The proof actually derives nonasymptotic (in *δ*) bounds on ϒ, but to simplify the exposition we choose to present the result in the form (45). To prove the theorem, we establish some lemmas that enable upper-bounding the distance between the probability measures corresponding to *H*_1_ and *H*_2_. For *i* = 1, 2, we suppose that under *H*_*i*_ the coalescent times are generated under a two-island model with rate matrix **Q**^(*i*)^ = *Q*(**c**^(*i*)^, **m**^(*i*)^). We assume that **Q**^(2)^ is *δ*-close to **Q**^(1)^. By rescaling coalescent time, we may assume ‖**Q**^(1)^‖ = 1, so that

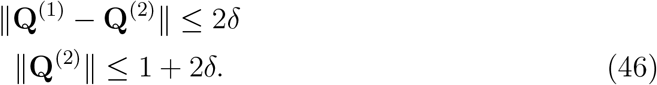

#### Lemma 16.

*Let* 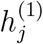 *and* 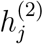 *be the hazard rate functions corresponding to H*_1_ *and H*_2_ *when there are j lineages remaining*.

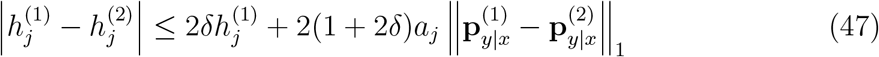

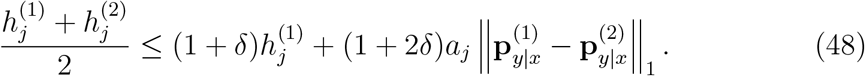

*Proof*. Letting 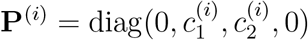 and recalling equation (13), we have

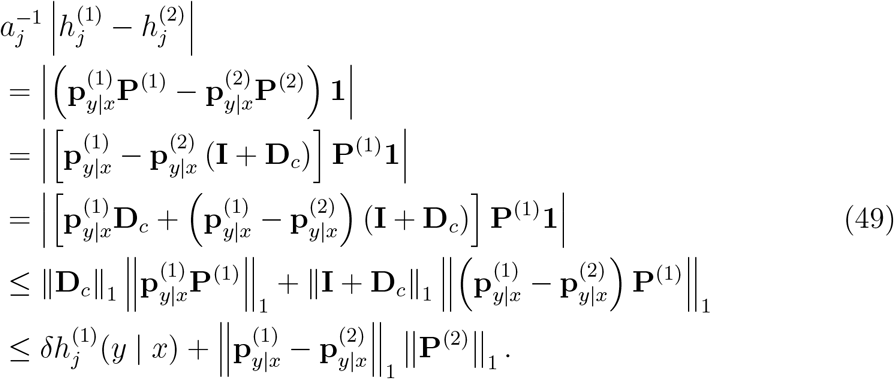

By equations (11) and (46),

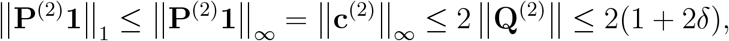

establishing (47). Inequality (48) follows by writing

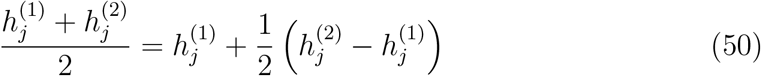

and using (47).

□

#### Lemma 17.

*Let* 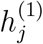 *and* 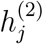 *as above. Then*

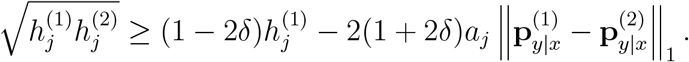

*Proof*. Using the identity

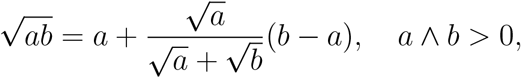

we have

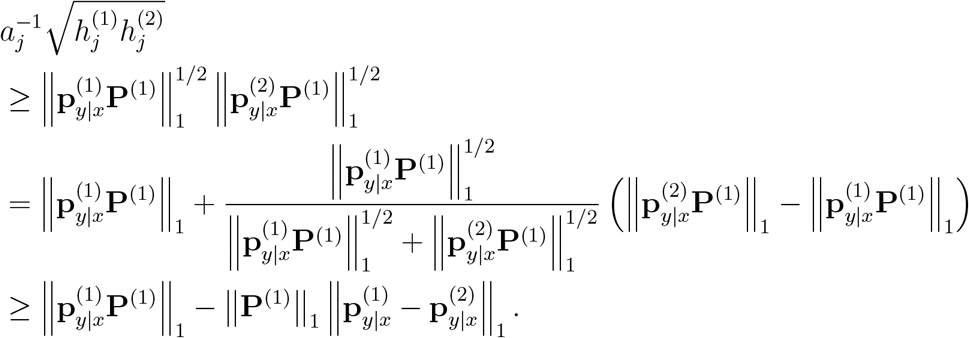

Multiplying both sides by *a*_*j*_ implies the result.

□

*Proof of Theorem 15*. The squared Hellinger distance between the two hypotheses is

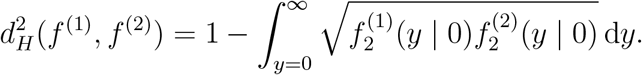

By Lemmas (16) and (17),

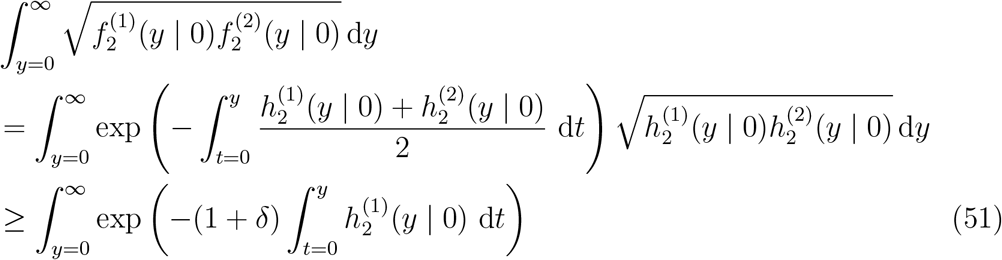

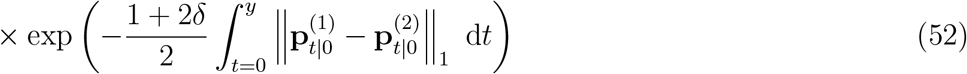

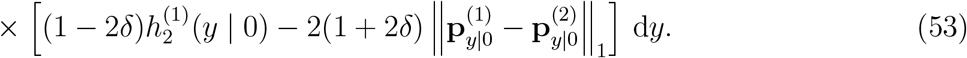

We combine lines (51) and (52) in the above display to form

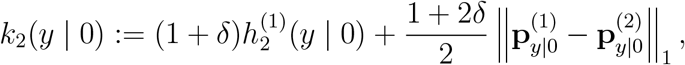

and then subtract (1 + *δ*)/(1 − 2*δ*) times line (53) from *k*_2_ to form

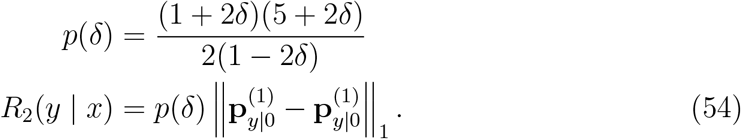

This gives us

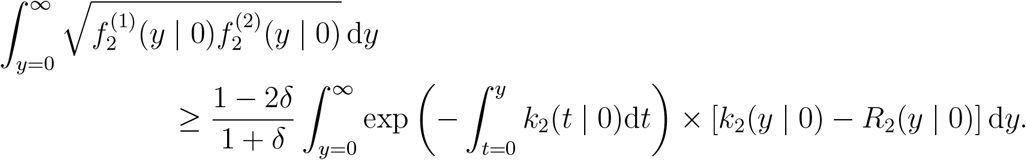

Splitting the integral into two pieces, we first have

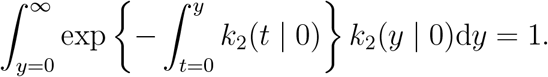

For the other piece, the Chernoff bound (12) gives

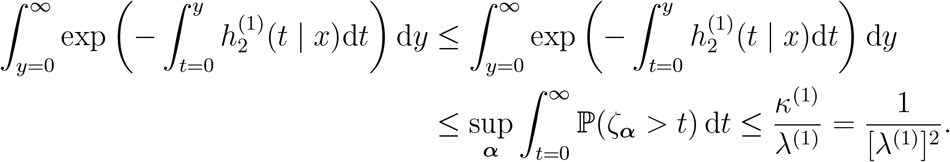

By Theorem 10,

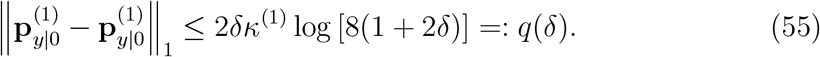

The preceding display and (54) imply

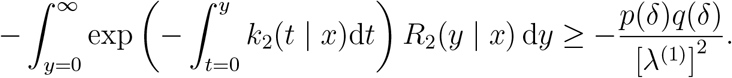

Putting it together, we have

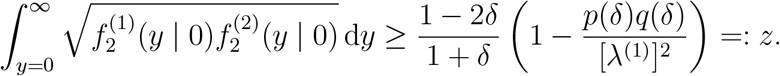

Expanding *z* in powers of *δ*, we find that

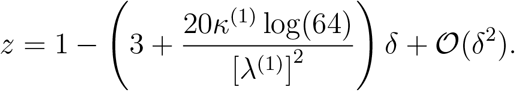

The Hellinger distance then satisfies

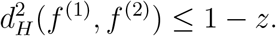

As *δ* tends to 0, we eventually have 0 *< z <* 1. Finally, we have 1 − *z*≤*𝒪* (*δ*), so we obtain (45).

□

*Remark*. In the proof above we used Theorem 10 to bound 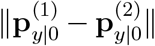 in equation (55). This makes use of the assumption that 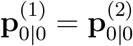, i.e. the starting distributions under the two hypotheses are the same. If we were to consider sample sizes larger than two, we would have to control the difference between the conditional distributions (see equation 3) 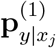 and 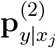, where *x*_*j*_ is a (random) time at which the *j*-th coalescent event takes place. This turns out to be difficult without placing additional and somewhat unnatural assumptions on **Q**^(1)^ and **Q**^(2)^, so the result is limited in its current form to the case *n* = 2. Note that this difficulty is specific to multi-population models and does not arise in the single-population analysis.

### 6.3 Isolation-with-migration

In this section, we consider a two-island problem where the two populations were part of a panmictic ancestral population until time *τ* > 0 in the past, sometimes referred to as the isolation-with-migration (IwM) model (Hey and Nielsen, 2007).

Let **c** = (*c*_0_, *c*_1_, *c*_2_) be the vector of coalescent rates where 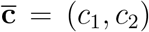 are for the islands under the two-island portion of the model and *c*_0_ is for the ancestral population. We will need

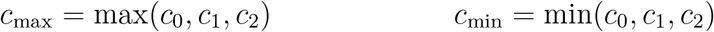

as well. As in the previous section, we consider the ability to distinguish between two hypothesized models, so 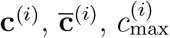, etc. are defined for *i* = 1, 2.

The hazard rate function now depends on *t* and *τ* :

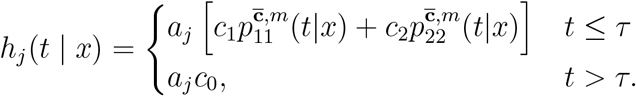

The final result is an analog of Theorem 15 for the case where two IwM models are compared.

#### Theorem 18.

*Let H*_1_ *and H*_2_ *be hypotheses with the same ancestral coalescent rate c*_0_ *and two-island rate matrix* **Q** *but different divergence times τ* ^(1)^ *and τ* ^(2)^ = (1+*δ*)*τ* ^(1)^. *Suppose L i*.*i*.*d. coalescent times on n individuals are sampled under H*_1_ *or H*_2_, *each with probability* 1/2, *then for sufficiently small δ* > 0, *the Bayes error rate for any classifier is at least* (1 − ϒ)/2, *where*

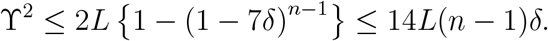

*Proof*. The proof is along the same lines as that of Theorem 15. Let 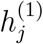 correspond to *τ* ^(1)^ and 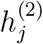 to *τ* ^(2)^. We have

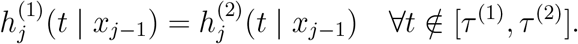

Otherwise,

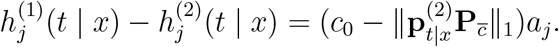

We quantify the difference in hazard rates by adding and subtracting terms. Let **1**(*τ* ^(1)^ ≤ *t* ≤ *τ* ^(2)^) be the indicator function that equals 1 whenever *τ* ^(1)^ ≤ *t* ≤ *τ* ^(2)^ and equals 0 otherwise. Then

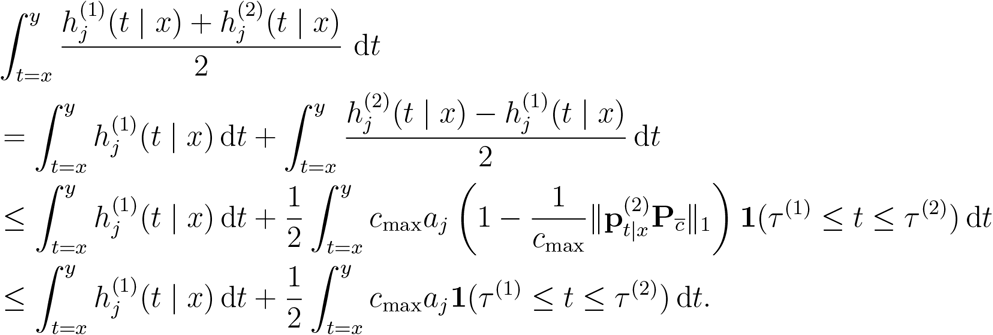

Now for some *α* ∈ [0, 1] we have

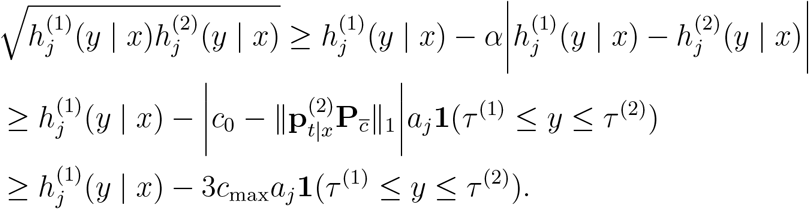

Combining the preceding displays, we get

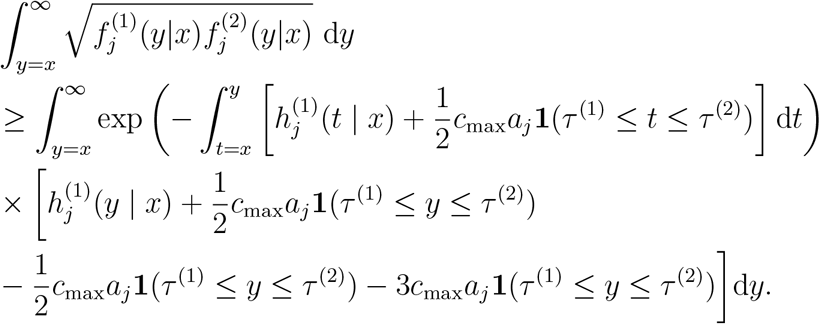

The first part integrates to one:

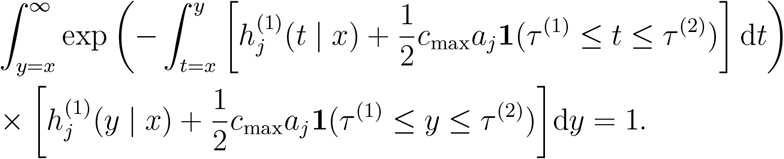

For the other part,

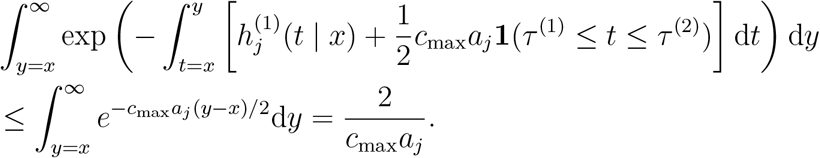

This implies

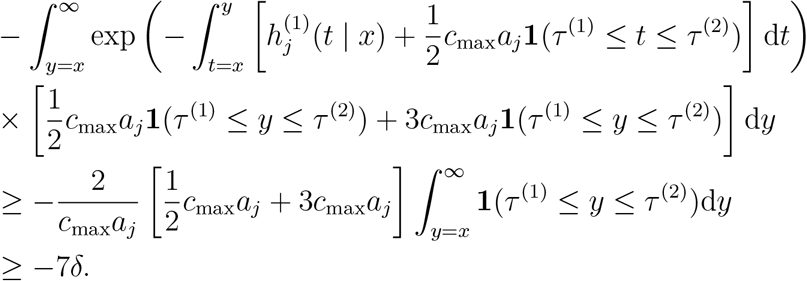

Putting it together, we have

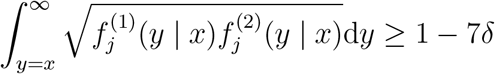

Hence,

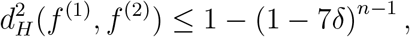

giving the first inequality. With *z* := 1 − 7*δ*, we use the bound 1 −*z*^*n−*1^ ≤ (*n −*1)(1 − *z*) to obtain the second inequality.

□

## 7 Discussion

In this paper, we studied upper and lower bounds for parameter estimation in the two-island model with migration. In Section 5 we derived some upper bounds on estimation error in the symmetric two-island model, and confirmed by simulations that our theoretical results are accurate (up to constant factors). In Section 6, we obtained lower bounds on the Bayes error rate for distinguishing between different two-island and isolation-with-migration models. Our results have basically the same consistent message: if the “sample size” *Ln* is much smaller than 1*/δ*, where *δ* is some measure of relative closeness between the hypotheses, then no procedure is able to reliably distinguish between them on the basis of sampled coalescent times.

It is instructive to compare our results to those of J. Kim et al., which inspired the present work and whose proof techniques we have adapted. Our results differ by leading order in *δ* (*η*, in their notation): J. Kim et al. obtain ϒ^2^ ≤ 𝒪 (*Lnδ*^2^) whereas the bounds in this paper are merely (at worst) 𝒪 (*Lnδ*). The bounds have different leading orders, and for small *δ* theirs is tighter. Note that their setting is not technically a special case of the one we consider here since we need to assume that **m** ≠ 0 in the definition of the model (2); if we do not assume this, the condition number *κ →* ∞ and many of the results in Section 4 become vacuous. At present, we do not know whether the difference is due to our proof method, or whether estimating the coalescent and migration rates may be easier under a multi-population model.

The models we have analyzed here are very basic, consisting of only a few parameters and at most two populations. Even if this restricted setting, the theoretical analysis is already cumbersome. Nowadays, significantly larger and more complicated models involving many populations and migration events between them are routinely estimated from large genetic datasets; there is a large gap between theory and practice. We have attempted to fill that gap, but there are many possible extensions and avenues for future work. In particular, we are not able to say anything about likelihood-based estimation in multi-population models, despite it being by far the dominant mode of method of estimation in applications. A useful, though seemingly difficult, future direction would be to study the likelihood function of genetic data under multi-population models with migration.

## Acknowledgments

BL is supported by NSF grant DMS-1646108. JT is supported by NSF grant DMS-2052653.

